# Genome-wide identification of binding sites of GRHL2 in luminal-like and basal A subtypes of breast cancer

**DOI:** 10.1101/2020.02.13.946947

**Authors:** Zi Wang, Haoyu Wu, Lucia Daxinger, Erik HJ Danen

**Author notes:** correspondence to Erik HJ Danen.

## Abstract

Grainyhead like 2 (GRHL2) is one of three mammalian homologues of the grainyhead (GRH) gene. It suppresses the oncogenic epithelial-mesenchymal transition (EMT), acting as a tumor suppressor. On the other hand, GHRL2 promotes cell proliferation by increasing human telomerase reverse transcriptase (hTERT) activity, serving as a tumor promoter. According to gene expression profiling, breast cancer can be divided into basal-like (basal A and basal B), luminal-like, HER2 enriched, claudin-low and normal-like subtypes. To identify common and subtype-specific genomic binding sites of GRHL2 in breast cancer, GRHL2 ChIP-seq was performed in three luminal-like and three basal A human breast cancer cell lines. Most binding sites of GRHL2 were found in intergenic and intron regions. 13,351 common binding sites were identified in basal A cells, which included 551 binding sites in gene promoter regions. For luminal-like cells, 6,527 common binding sites were identified, of which 208 binding sites were found in gene promoter regions. Basal A and luminal-like breast cancer cells shared 4711 GRHL2 binding sites, of which 171 binding sites were found in gene promoter regions. The identified GRHL2-binding motifs are all identical to a motif reported for human ovarian cancer, indicating conserved GRHL2 DNA-binding among human cancer cells. Notably, no binding sites of GRHL2 were detected in the promoter regions of several established EMT-related genes, including CDH1, ZEB1, ZEB2 and CDH2 genes. Collectively, this study provides a comprehensive overview of interactions of GRHL2 with DNA and lays the foundation for further understanding of common and subtype-specific signaling pathways regulated by GRHL2 in breast cancer.

## Introduction

Breast cancer is the predominant cause of cancer-related death in women aged 20 to 59 years globally [1]. Based on gene expression profiling, breast cancer can be divided into several subtypes with distinct molecular features, which includes luminal-like (luminal A and luminal B), basal-like (basal A and basal B), human epidermal growth factor receptor 2 (HER2)-enriched, claudin-low and normal-like subtypes[2]. Both luminal-like and basal-like subtypes comprise at least 73% of all breast cancers[2]. Conversions of luminal to basal lineage have been observed in mouse breast cancer models [3, 4] but luminal-like and basal-like subtypes differ in prognosis and response to therapy. Therefore, it is important to characterize common features and discordances between them.

The GRH gene was discovered in *Drosophila* and its mammalian homologs have three members (GRHL1, GRHL2 and GRHL3)[5]. GRH deficiency leads to failure of complete neural tube closure, epidermal barrier formation, trachea elongation and epidermal wound response [5–7]. GRHL2 is one of three mammalian homologues of the GRH gene, which has been investigated in cancer development. GRHL2 is located on chromosome 8q22 that is frequently amplified in many cancers, including breast cancer, colorectal cancer and oral squamous cell carcinoma[8–10]. GRHL2, as an oncogene, positively regulates cell proliferation by enhancing hTERT activity through inhibition of DNA methylation at 5’-CpG island around gene promoter[9]. GRHL2 inhibits cell apoptosis by suppressing death receptor (FAS and DR5) expression in breast cancer cells[8, 11]. Knockdown of GRHL2 downregulated ERBB3 expression, resulting in inhibition of cell proliferation[11]. On the other hand, GRHL2 was previously reported as a suppressor of oncogenic EMT by the loop of GRHL2-miR200-ZEB1 and regulation of the TGF-β pathway[12–14]. These controversial results suggest that the roles of GRHL2 may be tumor-specific through regulating different target genes in different cancers.

Chromatin immunoprecipitation followed by deep sequencing (ChIP-seq) is a widely used method to analyze protein-DNA interactions, histone modifications, and nucleosomes on genome-wide scale in living cells by capturing proteins at sites of their binding to DNA[15, 16]. Previous findings showed that GRHL2 shares a similar DNA-binding motif with other GRHL family members[13, 17, 18]. To date, no studies have investigated the genomic landscape of GRHL2 binding sites across breast cancer subtypes. In this study, we provide a comprehensive overview of binding sites of GRHL2 in the genome of basal A and luminal-like subtypes of breast cancer.

## Methods and materials

### Cell lines

Human breast cancer cell lines representing luminal-like (MCF7, T47D, BT474), basal A (HCC1806, BT20 and MDA-MB-468), and basal B subtypes (Hs578T) were obtained from the American Type Culture Collection. Cells were cultured in RPMI1640 medium with 10% fetal bovine serum, 25 U/mL penicillin and 25 μg/mL streptomycin in the incubator (37°C, 5% CO2).

### Chromatin immunoprecipitation-sequencing (ChIP-seq)

Cells were grown in RPMI-1640 complete medium. Cross-linking was performed by 1% formaldehyde for 10 minutes at room temperature (RT). Then 1M glycine (141 μl of 1M glycine for 1 ml of medium) was used to quench for 5 minutes at RT. Cells were washed twice with icecold PBS containing 5 μl/ml phenylmethylsulfonyl fluoride (PMSF). Cells were harvested by centrifugation (2095 g for 5 minutes at 4°C) and lysed with NP40 buffer (150 mM NaCl, 50mM Tris-HCl, 5mM EDTA, 0.5% NP40, 1% Triton X-100) containing 0.1% SDS, 0.5% sodium deoxycholate and protease inhibitor cocktail (EDTA-free Protease Inhibitor Cocktail, Sigma). Chromatin was sonicated to an average size of 300 bp **(Fig. S1)**. GRHL2-bound chromatin fragments were immunoprecipitated with anti-GRHL2 antibody (Sigma; HPA004820). Precipitates were eluted by NP buffer, low salt (0.1% SDS, 1% Triton X-100, 2mM EDTA, 20mM Tris-HCl (pH 8.1), 150mM NaCl), high salt (0.1% SDS, 1% Triton X-100, 2mM EDTA, 20mM Tris-HCl (pH 8.1), 500mM NaCl) and LiCl buffer (0.25M LiCl, 1%NP40, 1% deoxycholate, 1mM EDTA, 10mM Tris-HCl (pH 8.1)). Chromatin was de-crosslinked by 1% SDS at 65°C. DNA was purified by Phenol:Chloroform:Isoamyl Alcohol (PCI) and then diluted in TE buffer.

In order to examine the quality of our samples before sequencing, ChIP-PCR was performed to validate interaction of GRHL2 with the promoter region of CLDN4, a direct target gene GRHL2 [19]. The results confirmed the GRHL2 binding site around the CLDN4 promoter **(Fig. S2)**. The following primers were used for ChIP-PCR: CLDN4 forward: gtgacctcagcatgggctttga, CLDN4 reverse: ctcctcctgaccagtttctctg, Control (an intergenic region upstream of the GAPDH locus) forward: atgggtgccactggggatct, Control reverse: tgccaaagcctaggggaaga. ChIP-PCR data were collected and analyzed using the 2^-ΔΔCt^ method [20].

For ChIP-Seq, library preparation and paired-end sequencing were performed by GenomeScan (Leiden, The Netherlands)

### ChlP-seq data analysis

Paired-end reads were mapped to the human reference genome (hg38) using BWA-MEM[21] with default parameters. Over 93% of total reads were mapped to the human genome in BT20, HCC1806, MDA-MB-468, T47D and MCF7 cell line. For BT474, ~57.3% reads were mapped. Phred quality score (Q score) was used to measure base calling accuracy, which indicates the probability that a given base is called incorrectly[22, 23]. Q score is logarithmically related to the base calling error probabilities *P* [22, 23].

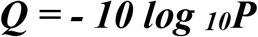

Q=30 nominally corresponds to a 0.1% error rate[24]. Reads with scores > Q30 were over 86% in BT20, HCC1806, MDA-MB-468, T47D and MCF7 cell lines. For BT474, reads with scores > Q30 accounted for 48.6%.

To examine whether the paired-end reads were appended with unwanted adapter sequences, an adapter content test was performed. The quality control report **(Fig. S3)** showed that cumulative presence of adapter sequences was <5% in all cell samples, indicating that all data sets could be further analyzed without adapter-trimming. Per base sequence quality of sequencing was examined, which indicated that all sequencing data were of high quality **(Fig. S4)** and could be further analyzed.

Reads with low mapping quality (≤ Q30) were filtered out. MACS version 2.1.0[25] was used for peak calling by default settings. q value was adjusted to 0.1 for BT474 cell line to avoid loss of peaks. The annotatePeaks and MergePeaks function from HOMER[26] were used to annotate and overlap peaks, respectively. Motif analysis was performed by ChIP-seq peaks with high scores using the MEME-ChIP program with default settings.

## Results

### Genome-wide identification of binding sites of GRHL2 in luminal-like and basal A subtypes of breast cancer

To identify GRHL2 binding sites, ChIP-seq was performed in luminal-like (MCF7, T47D and BT474) and basal A (HCC1806, BT20 and MDA-MB-468) breast cancer cells. Firstly, the coverage of peak regions across chromosomes was analyzed[27]. In each cell sample, GRHL2 was strongly associated with all chromosomes **(Fig. 1)**, except for chromosome Y. The number of peaks on chromosome Y was minimal, partially due to the fact that Y chromosome did not have as many genes as other chromosomes had.

**Fig. 1.**
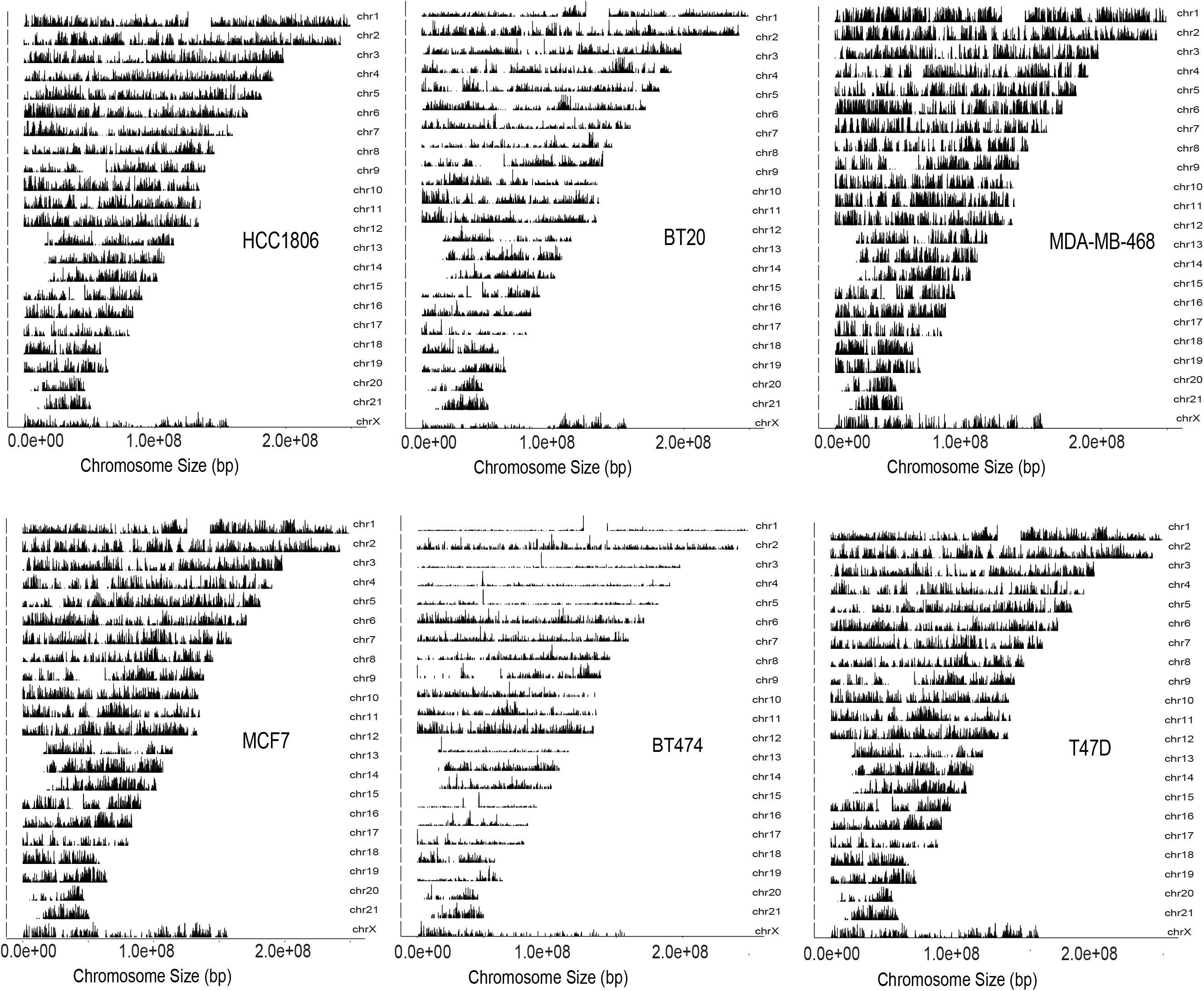
Coverage of peak regions across chromosomes. The graph represents the coverage of GRHL2 binding sites across the chromosomes.

GRHL2 binding sites were found in intergenic regions, transcription start sites (TSS) promoter regions, introns, exons, transcription termination sites (TTS) and unknown regions **(Fig. 2)**. The majority of peaks was located in intergenic and intron regions in basal A and luminal-like breast cancer cells. Genes where GRHL2 was found to interact with the −1000 to +100 promoter region in all three luminal (left column), all three basal A (middle column), or all luminal and basal A cell lines tested (right column) were identified and represent likely candidate general and subtype-specific GRHL2 target genes (Table S1).

**Fig. 2.**
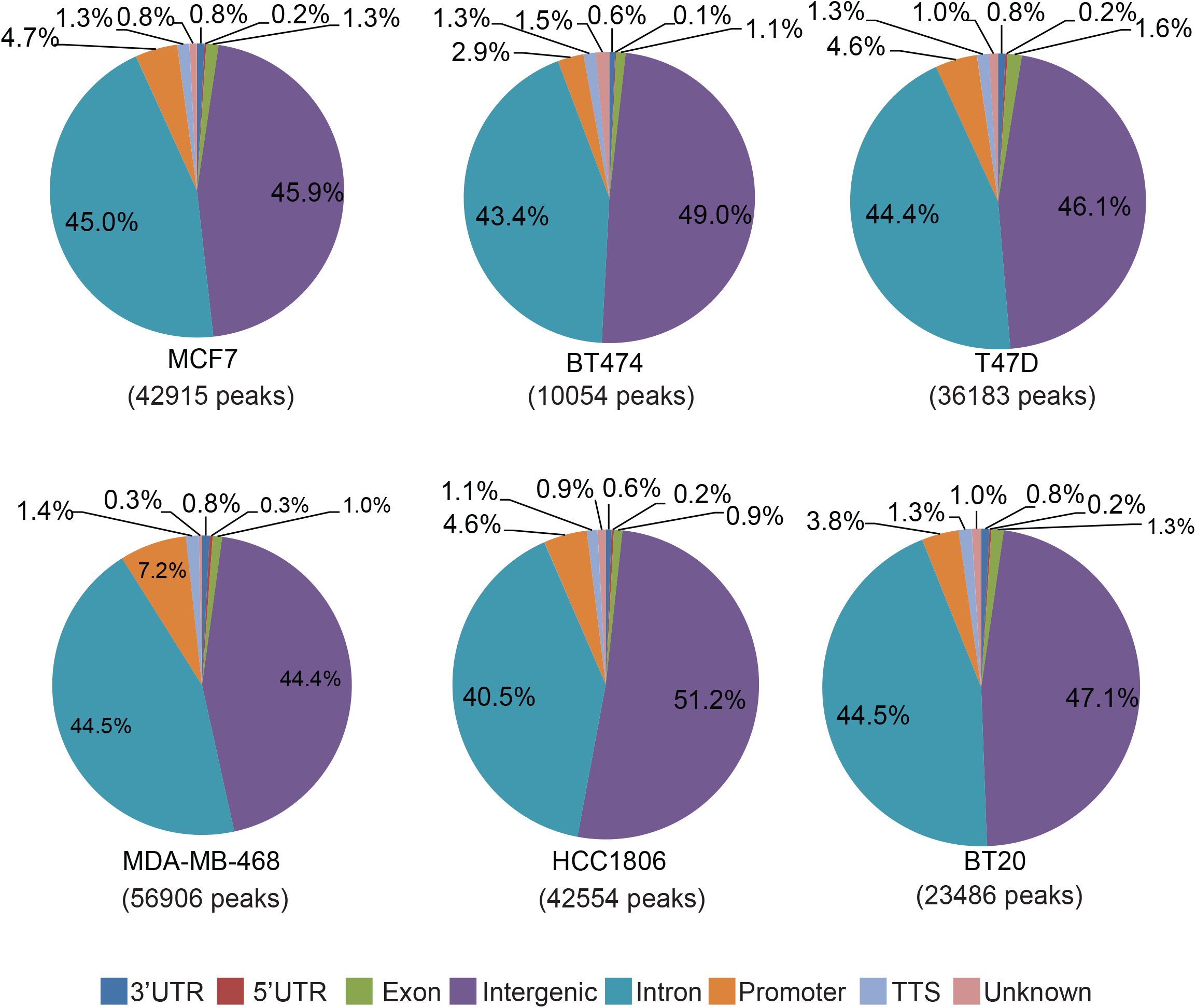
Percentage of GRHL2 binding sites found at promoter regions, 5’ untranslated regions (UTRs), 3’ UTRs, exons, introns, intergenic regions, transcription termination sites (TTSs) and unknown regions. Promoter regions are defined as −1000 bp to 100 bp from the transcription start sites (TSS).

To further investigate if peaks were enriched in promoter regions, read count frequency and density profiling of GRHL2 binding sites within −6000bp ~ +6000bp of the transcription start site (TSS) were analyzed **(Fig. 3)**. Consistent with the annotation of binding sites, which showed most GRHL2 binding sites existed in the intergenic regions, the density of GRHL2 binding sites was not increased in the −1000 to +100 promoter region of basal A and luminal-like breast cancer cells.

**Fig. 3.**
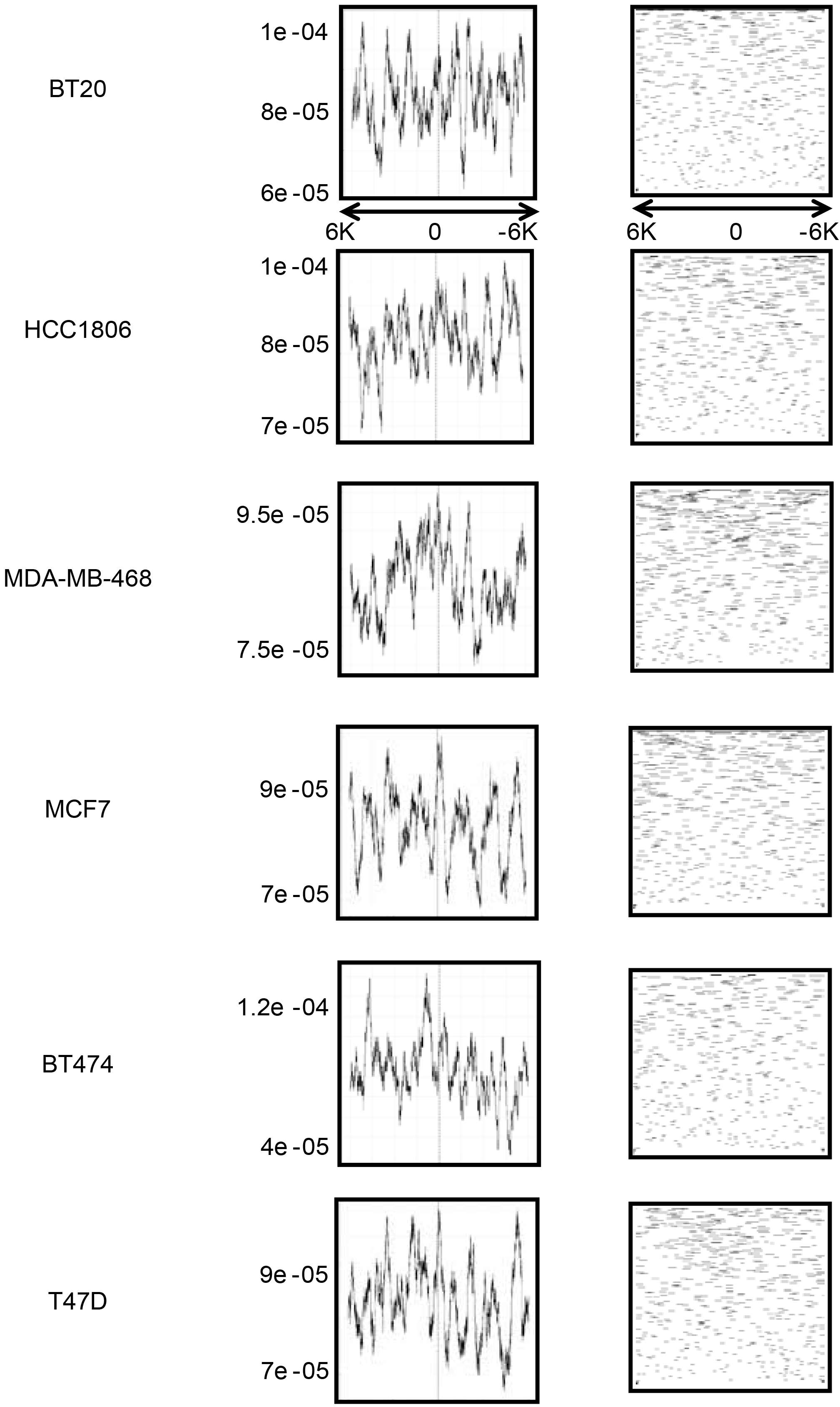
The read count frequency and density profile of GRHL2 binding sites within −6000 bp~ +6000 bp of the promoter-TSSs. On the left side, graphs are for GRHL2 ChIP-seq read count frequency in indicated cell line. X axis represents read count frequency; Y axis is for genomic region. On the right side, graphs show the density of ChIP-seq reads for GRHL2 binding sites in the indicated cell line.

To detect similarities of GRHL2 binding sites between luminal-like and basal A subtype, three luminal-like/basal A data sets were overlapped to identify shared binding sites. 13,351 common binding sites were identified in basal A subtype of breast cancer cells, which included 551 binding sites in gene promoter regions (−1000 ~ +100 from TSS) **(Fig. 4A and B)**. For luminal-like breast cancer cells, 6,527 common binding sites were identified, of which 208 binding sites were found in gene promoter regions **(Fig. 4C and D)**. Basal A and luminal-like subtypes of breast cancer cells shared 4,711 binding sites of GRHL2, of which 171 binding sites were found in gene promoter regions **(Fig. 4E and F)**.

**Fig. 4.**
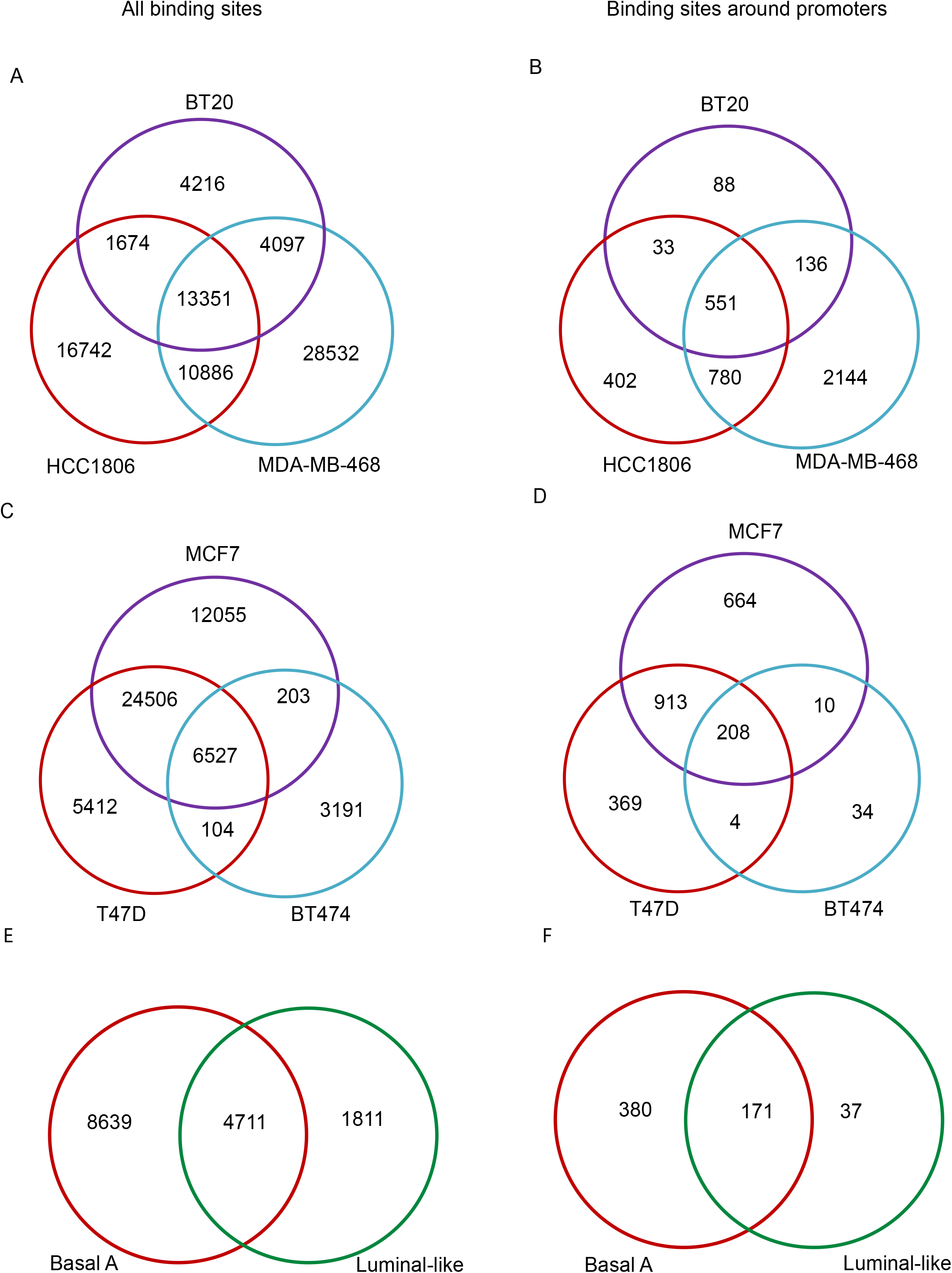
Overlap of GRHL2 binding sites. Overlap of GRHL2 binding sites is identified in the indicated subtypes.

### Identification of a common GRHL2-interaction motif

The MEME-ChIP program was used to identify motifs, all of which were with statistical significance. In each sample, 3 motifs with high E value were shown **(Fig. 5)**, whose core binding was similar to previously published ones [13, 28–30]. Thus, our ChIP-seq data indicated that GRHL2 motif was highly conserved in human and mouse cells.

**Fig. 5.**
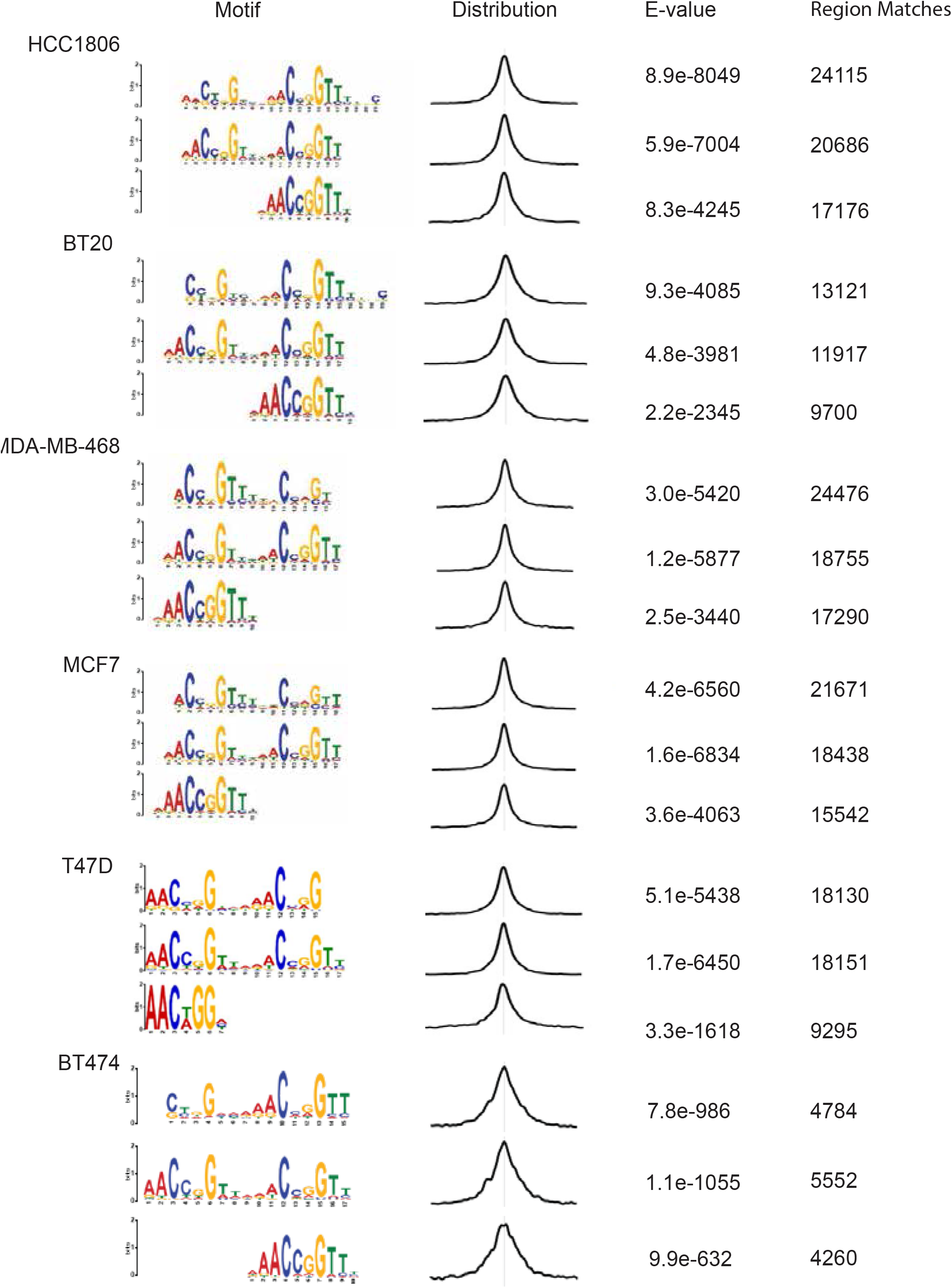
DNA-binding motif of GRHL2 in luminal-like and basal A subtypes of breast cancer. From left to right, the first panel shows the identified motifs in the indicated cells. The second panel shows distribution of the best matches to the motif in the sequences. The third panel shows E-value, the significance of the motif according to the motif discovery. The last panel shows the number of regions that match the corresponding motif.

### GRHL2-binding at EMT-related genes

GRHL2 and OVOL2 support an epithelial phenotype and counteract EMT transcription factor such as ZEB and SNAIL. Some studies have reported that GRHL2 binding sites are present in the intronic region of CDH1 and in the promoter regions of CLDN4 and OVOL2 for activation of transcription and GRHL2 was found to bind the ZEB1 gene as a negative regulator[12, 28, 31, 32]. In our ChIP-seq data, GRHL2 binding sites were observed at CDH1 introns and at promoter regions of CLDN4 and OVOL2 (Fig. 6). In our study, CLDN4 showed multiple GRHL2 binding sites across the gene coding/non-coding regions, suggesting the binding of GRHL2 to multiple regions may be involved in long-distance chromatin interactions as suggested previously[13]. Conversely, no GRHL2 binding was observed at the promoter of ZEB1 and ZEB2 (Fig. 6), arguing against mutual regulation through direct interaction as previously suggested [33]. These findings suggest that GRHL2 binding sites in EMT-related genes may be cell context-dependent.

**Fig. 6.**
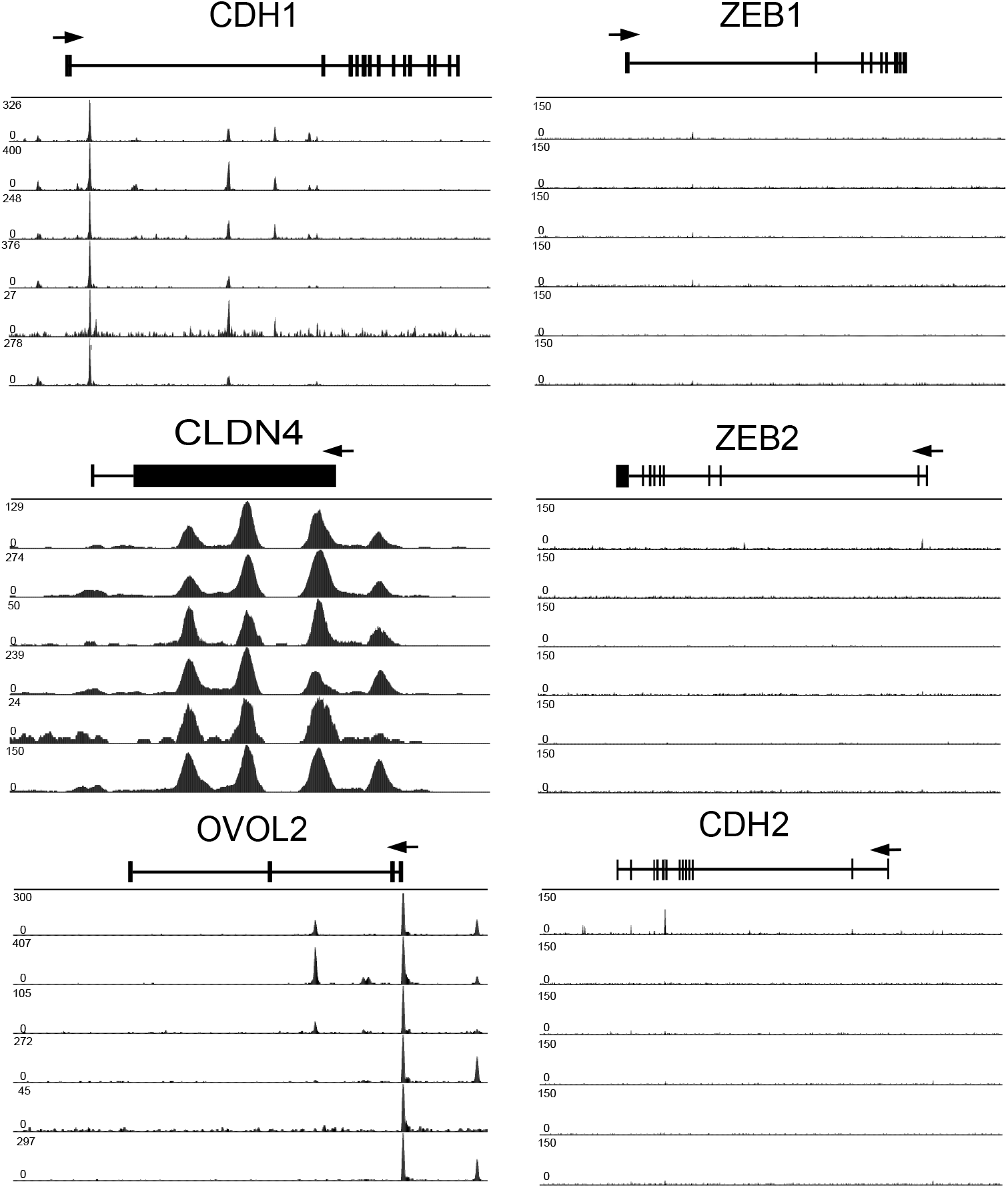
GRHL2 ChIP tracks at selected core genes and EMT genes. ChIP tracks are shown from top to bottom for HCC1806, MDA-MB-468, BT20, MCF7, BT474 and T47D, respectively. The snapshot on the left shows results for 3 identified GRHL2 targets (CDH1, CLDN4 and OVOL2) and the snapshot on the right shows results for three not-identified genes encoding proteins associated with EMT (ZEB1, ZEB2 and TWIST2). The track height is scaled from 0 to the indicated number. Above all tracks, the locus with its exon/intron structure is presented.

## Discussion

Cell type origin is one of the most important factors that determine molecular features of tumors[34]. In general, luminal-like tumor cells are biologically similar to cells derived from inner (luminal) cells lining the mammary ducts, whereas cells of basal-like breast cancer are characterized by features similar to surrounding the mammary ducts[35]. Basal-like breast cancers are associated with a worse prognosis and an increased possibility of cancer metastasis compared with the luminal-like subtype[4, 36, 37]. Immunohistochemical staining is clinically used to categorize luminal-like breast cancer into luminal A (estrogen receptor (ER) and/or progesterone receptor (PR) positive, HER2 negative) and luminal B (ER and/or PR, and HER2 positive). However, most basal-like breast cancers are negative for ER, PR and HER2, therefore the majority of basal-like breast cancer is triple negative breast cancer (TNBC). Basal-like breast cancer can be further subdivided into basal A and basal B. As for basal A, it is associated with BRCA1 signatures and resembles basal-like tumors, whereas basal B subtype displays mesenchymal properties and stem/progenitor characteristics[38, 39].

In the present study, ChIP-seq was performed to characterize genome-wide binding sites of transcription factor GRHL2 in basal A and luminal-like subtypes of breast cancer. The match with previously a published binding motif shows that GRHL2-interaction with the DNA is highly conserved in human cancer cells. A limited number of binding sites were located in gene promoter regions. Similar to previous reports [13, 29], most binding sites were located in introns and intergenic regions of target genes. Such regions may contain enhancers interacting with GRHL2 and GRHL2 has also been reported to regulate histone modifications such as H3K4me3 and H3K4me1[13, 40]. Together, this suggests that GRHL2 may regulate gene expression through direct transcriptional control at promoter regions or through alternative mechanisms including epigenetic mechnisms.

Close to 5000 identified GRHL2 genomic binding sites were shared between all tested basal A and luminal-like cell lines. A similar number of binding sites were found in all basal-like cell lines but were not detected in any of the tested luminal lines. These candidate subtype-specific GRHL2-target sites may serve as a starting point to the unraveling of distinct transcriptional networks in different breast cancer subtypes.

Our analysis of GRHL2 interaction with known EMT-related genes fits previously published findings except for ZEB1. It was reported that ZEB1 is regulated by GRHL2 directly and, vice versa, that ZEB1 regulates GRHL2 in a balance between EMT and MET [10–12, 33]. However, we did not detect obvious GRHL2 binding sites in the promoter regions of the ZEB1 or ZEB2 genes. GRHL2 may regulate ZEB1 and ZEB2 indirectly in luminal-like and basal A breast cancers.

Taken together, this study provides a comprehensive genome-wide resource of GRHL2 binding sites and identifies specific and shared binding sites for GRHL2 in luminal-like and basal A subtype breast cancer. Overall, this study lays the foundation for unraveling signaling pathways regulated by GRHL2.

## Supplemental data

**Table S1.**
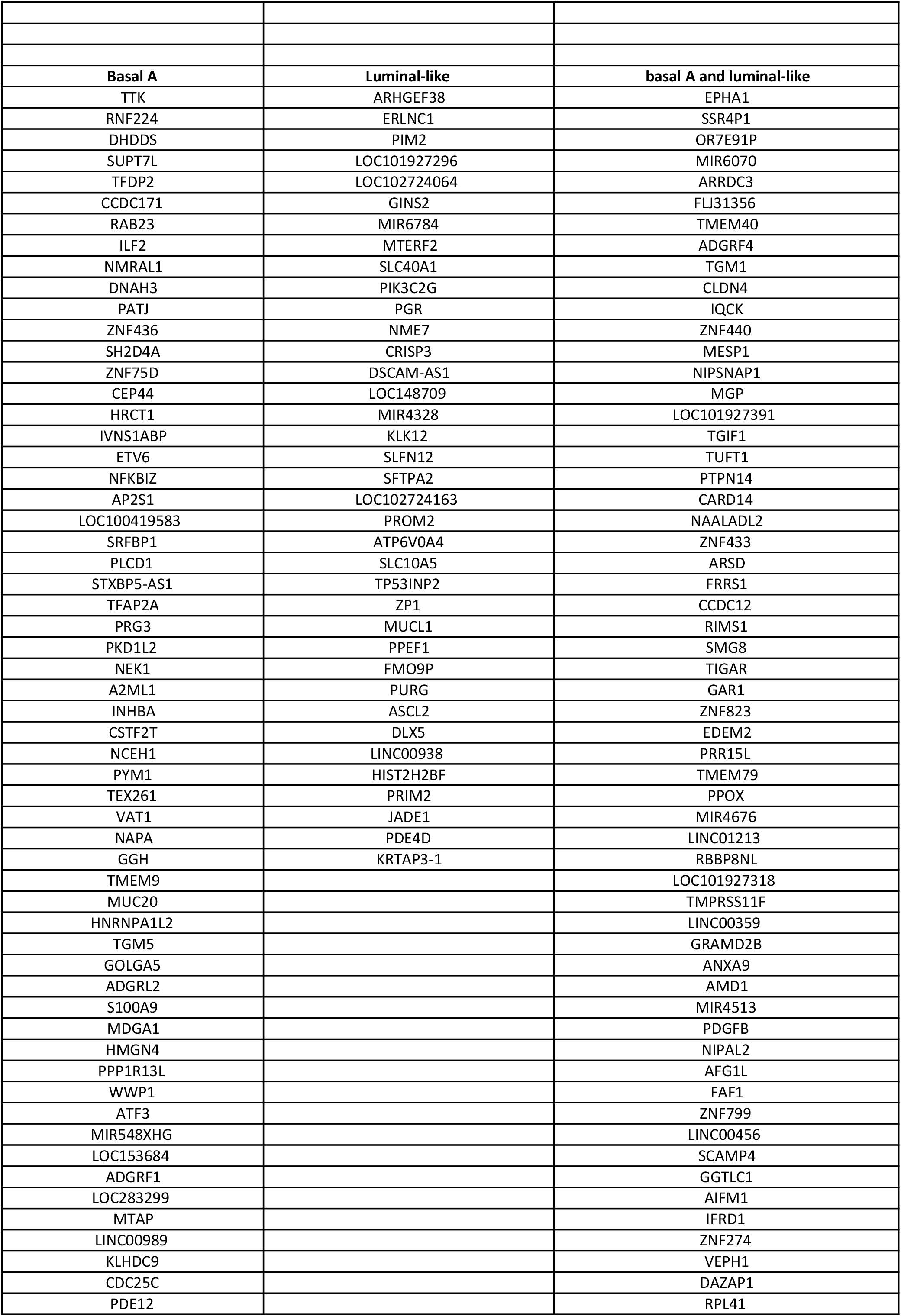
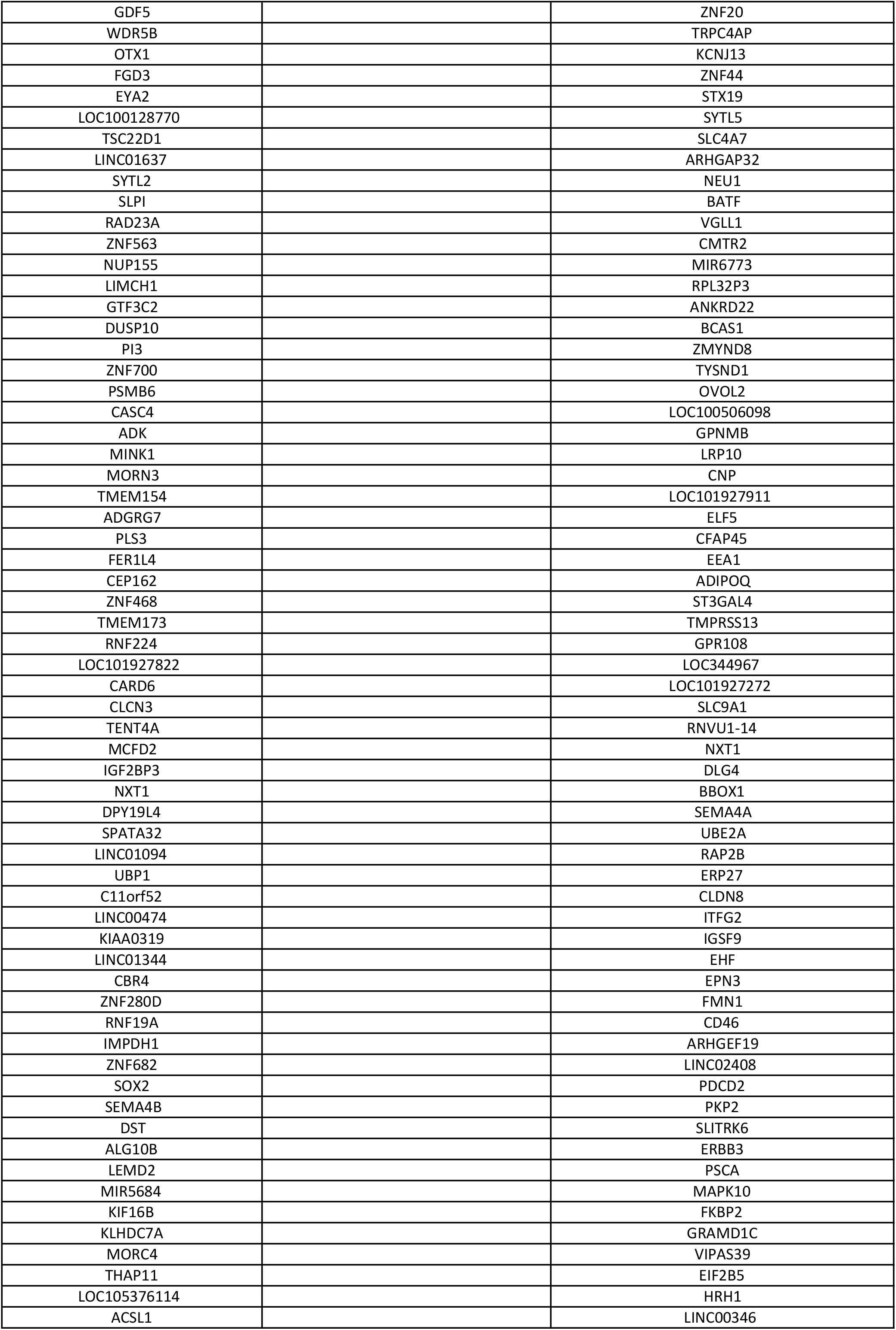
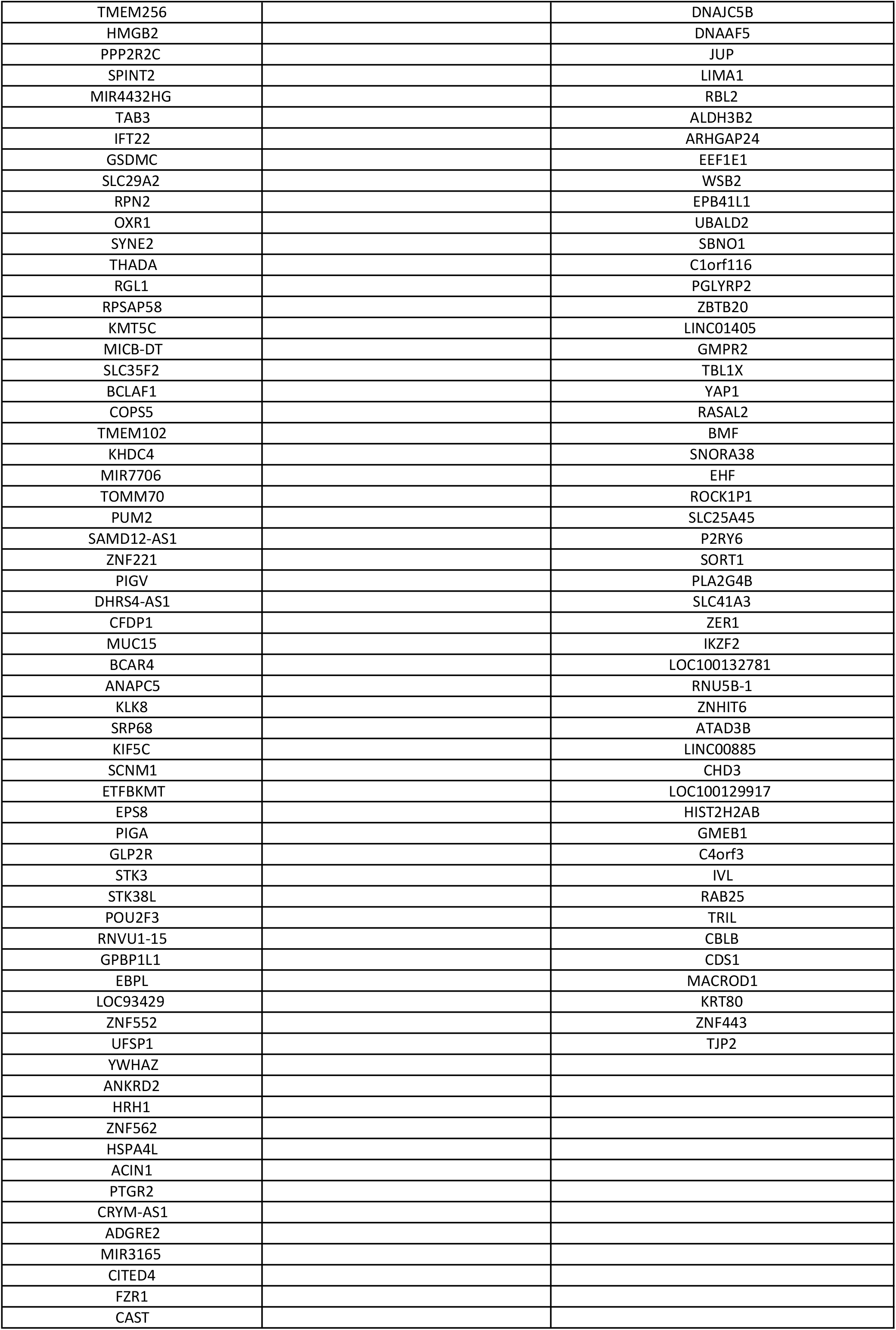
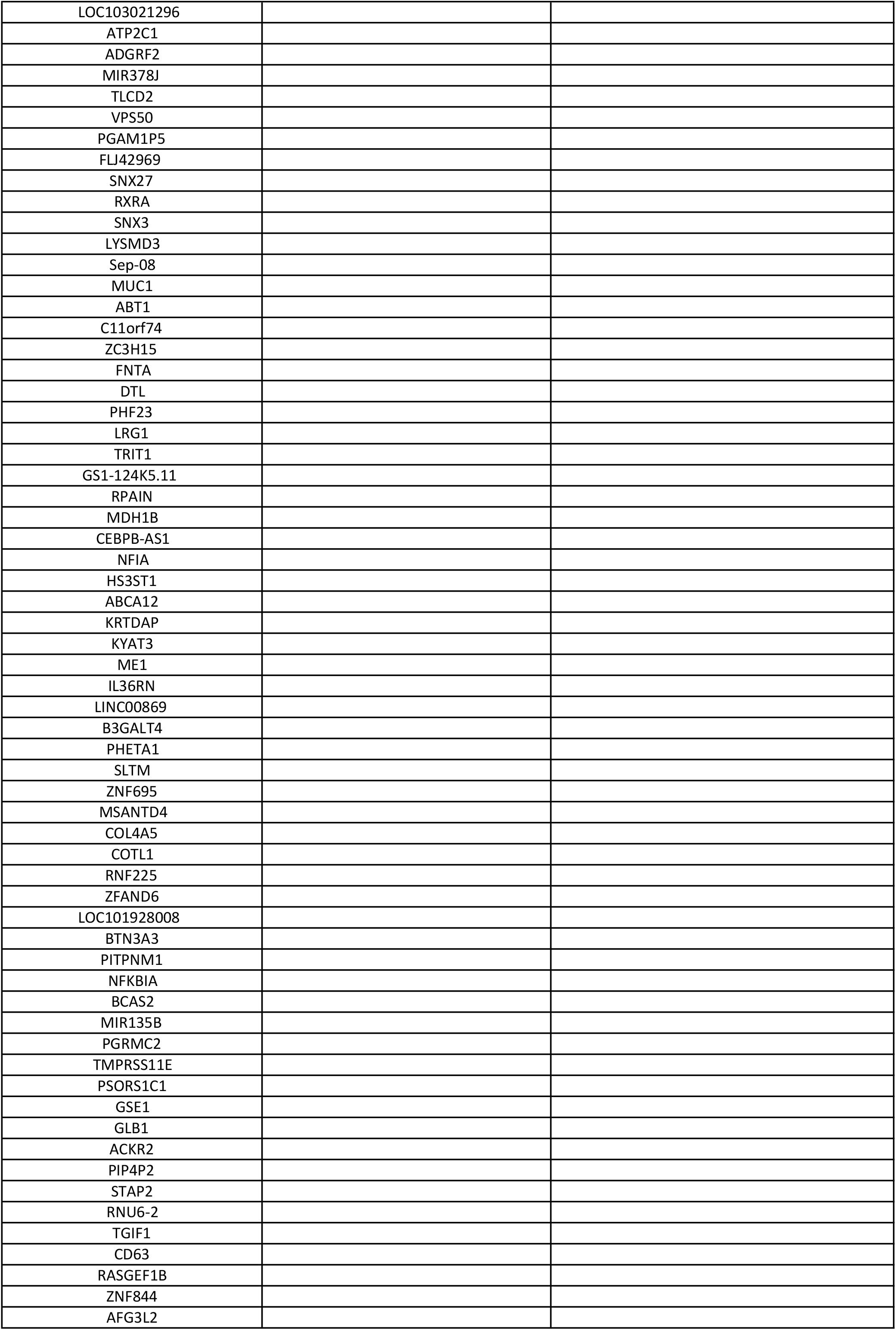
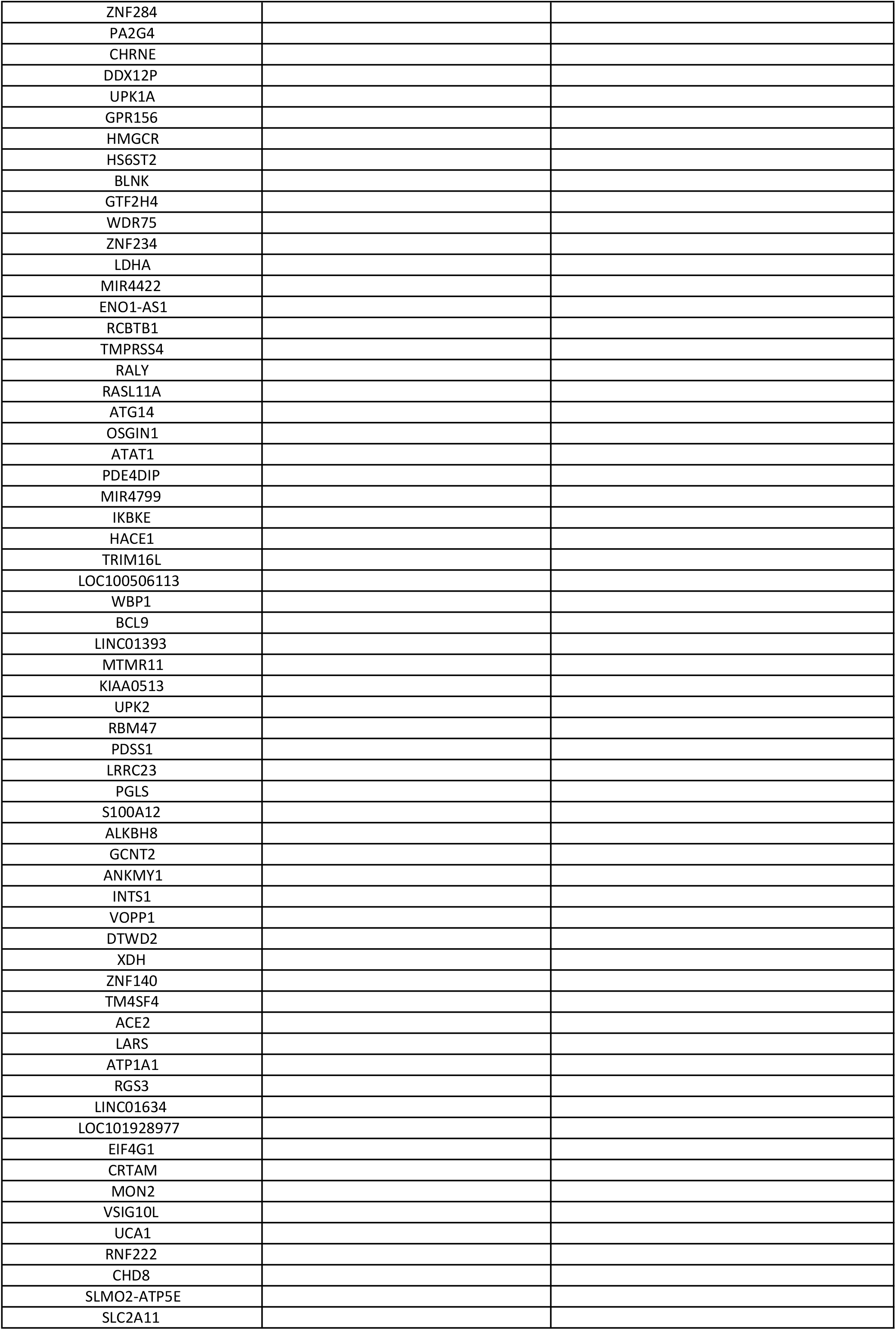
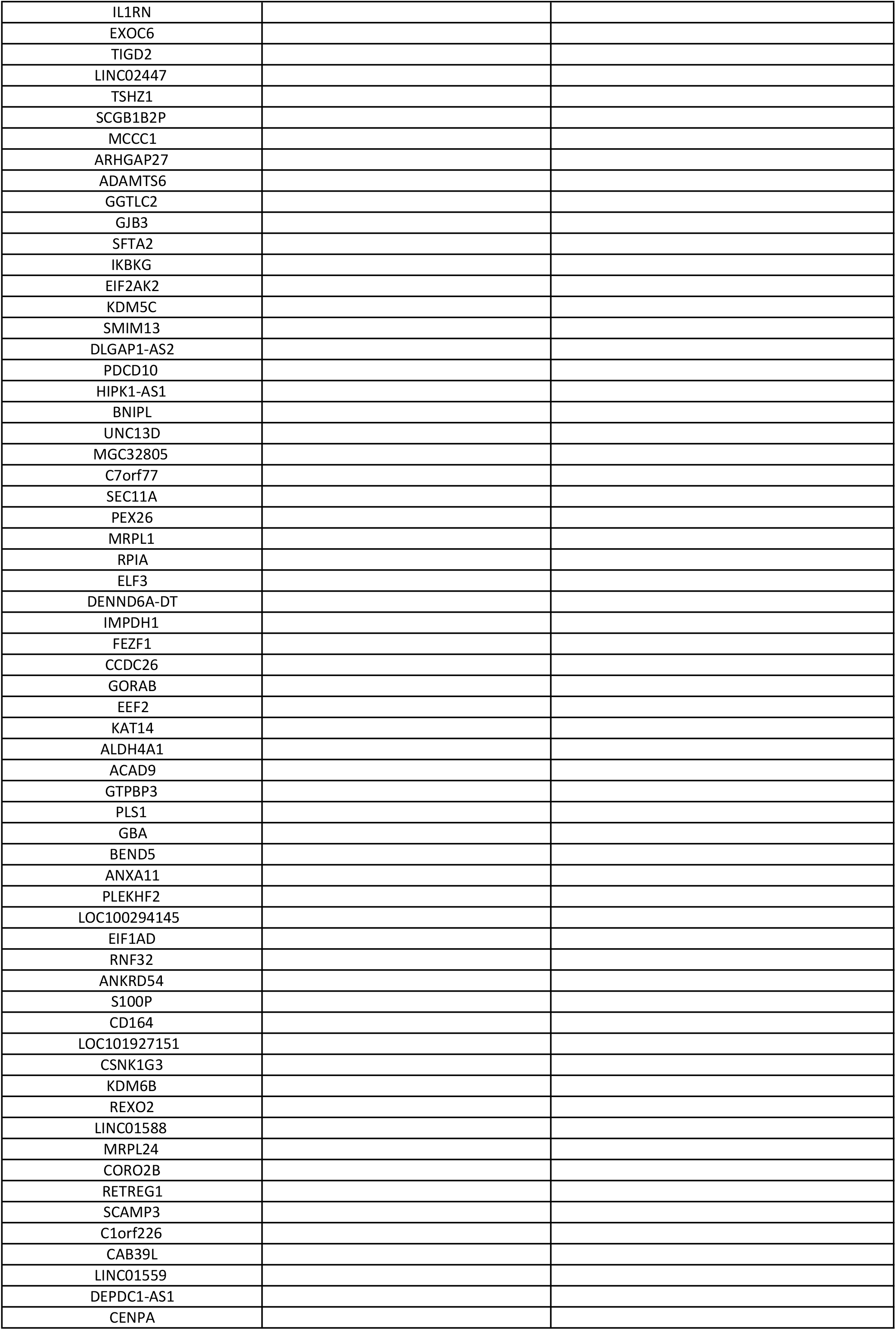
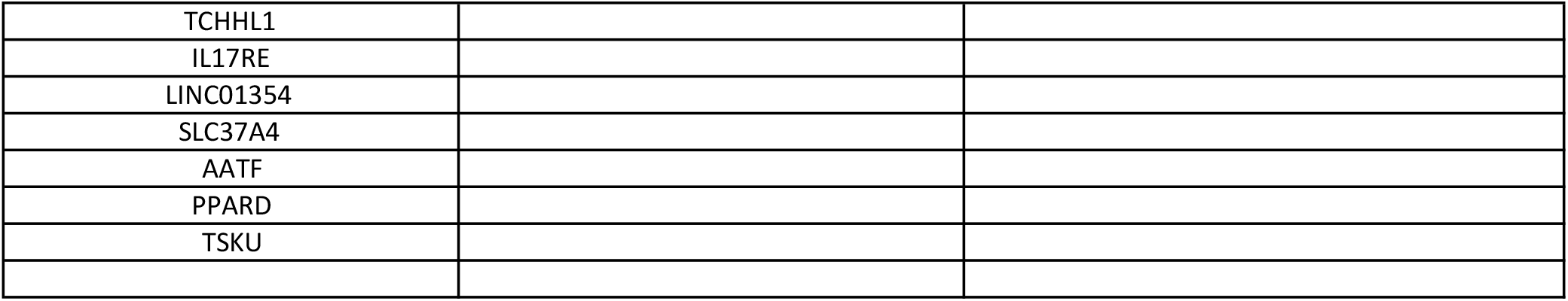
Candidate GRHL2 target genes. List of genes where GRHL2 was found to interact with the −1000 to +100 promoter region in all three basal A (left column), all three luminal-like (middle column), or all basal A and luminal-like cell lines tested (right column).

**Fig. S1.**
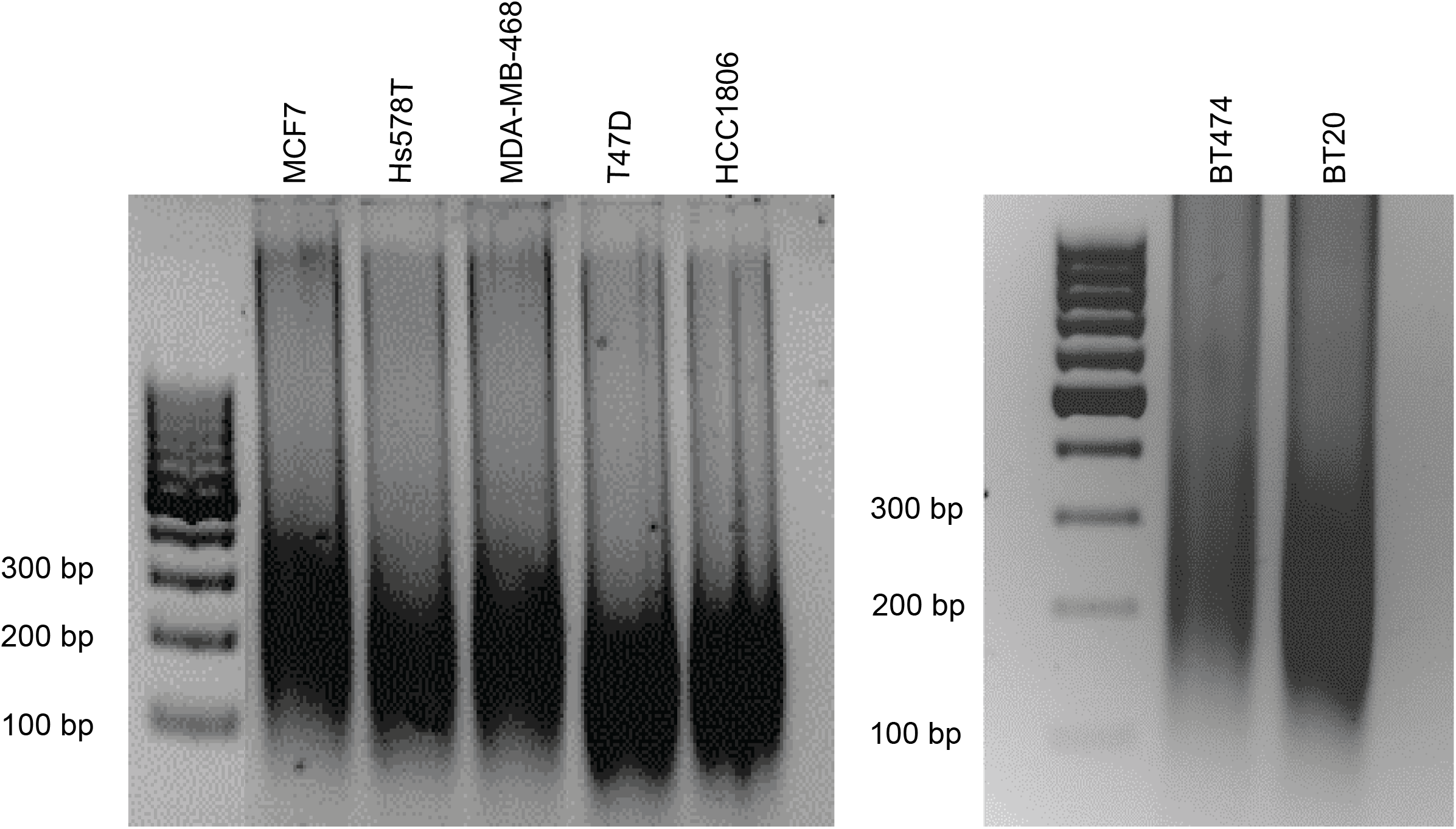
DNA fragmentation analysis by agarose gel electrophoresis. After sonication, indicated samples were purified and loaded to 2% agarose gel.

**Fig. S2.**
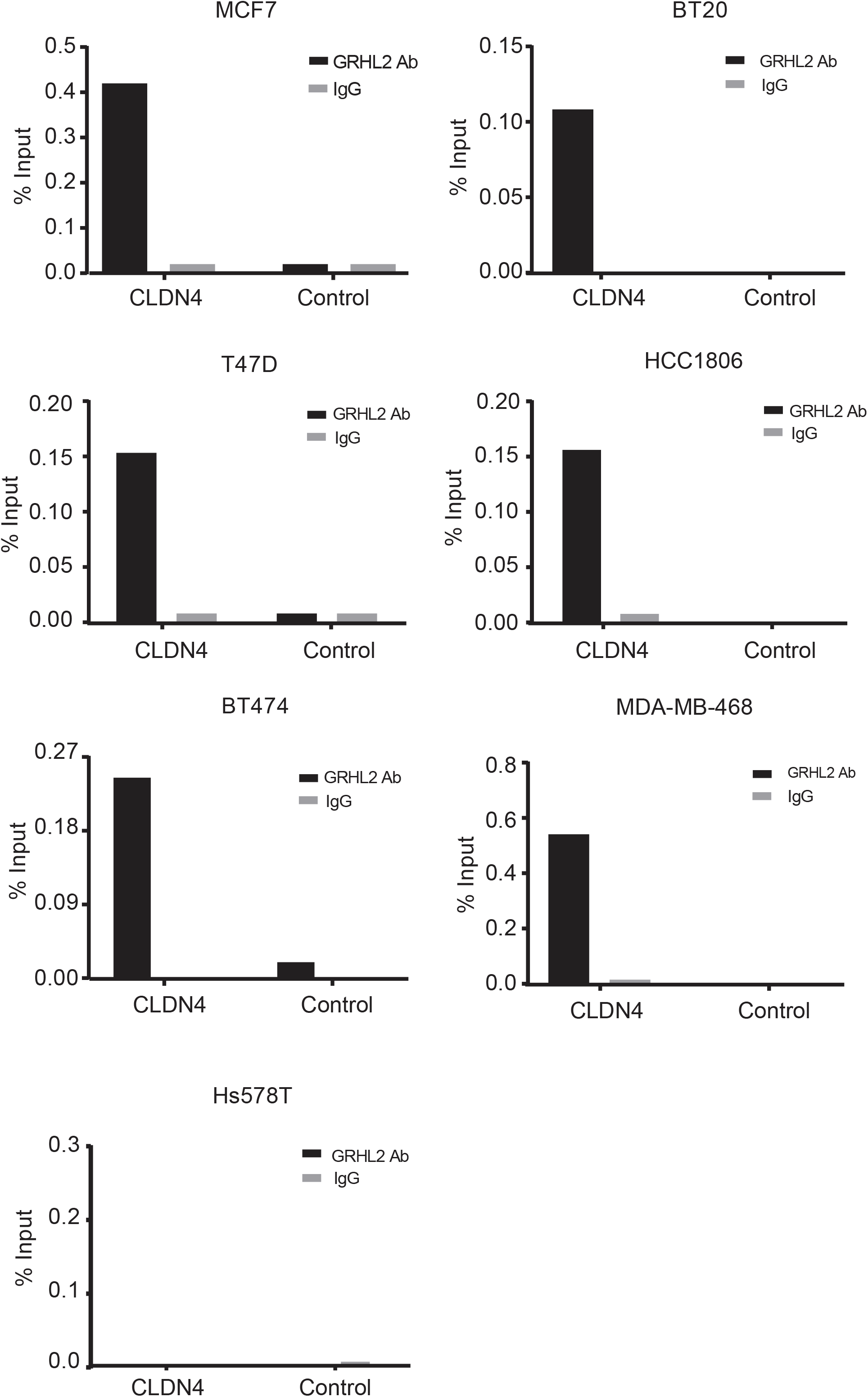
ChIP-PCR validation of the isolated genomic DNA fragments. Graphs represent the efficiency of CLDN4 genomic DNA co-precipitation with anti-GRHL2 Ab (black bars) or IgG control Ab (grey bars). Detection was performed by PCR using primers targeting the promoter region of CLDN4 or targeting the intergenic region upstream of the GAPDH locus (Control). Results are shown for 3 GRHL2-positive luminal cell lines (MCF7, BT474, T47D), 3 GRHL2-positive basal-A cell lines (BT20, HCC1806, MDA-MB-468), and 1 GRHL2-negative basal-B cell line (Hs578T).

**Fig. S3.**
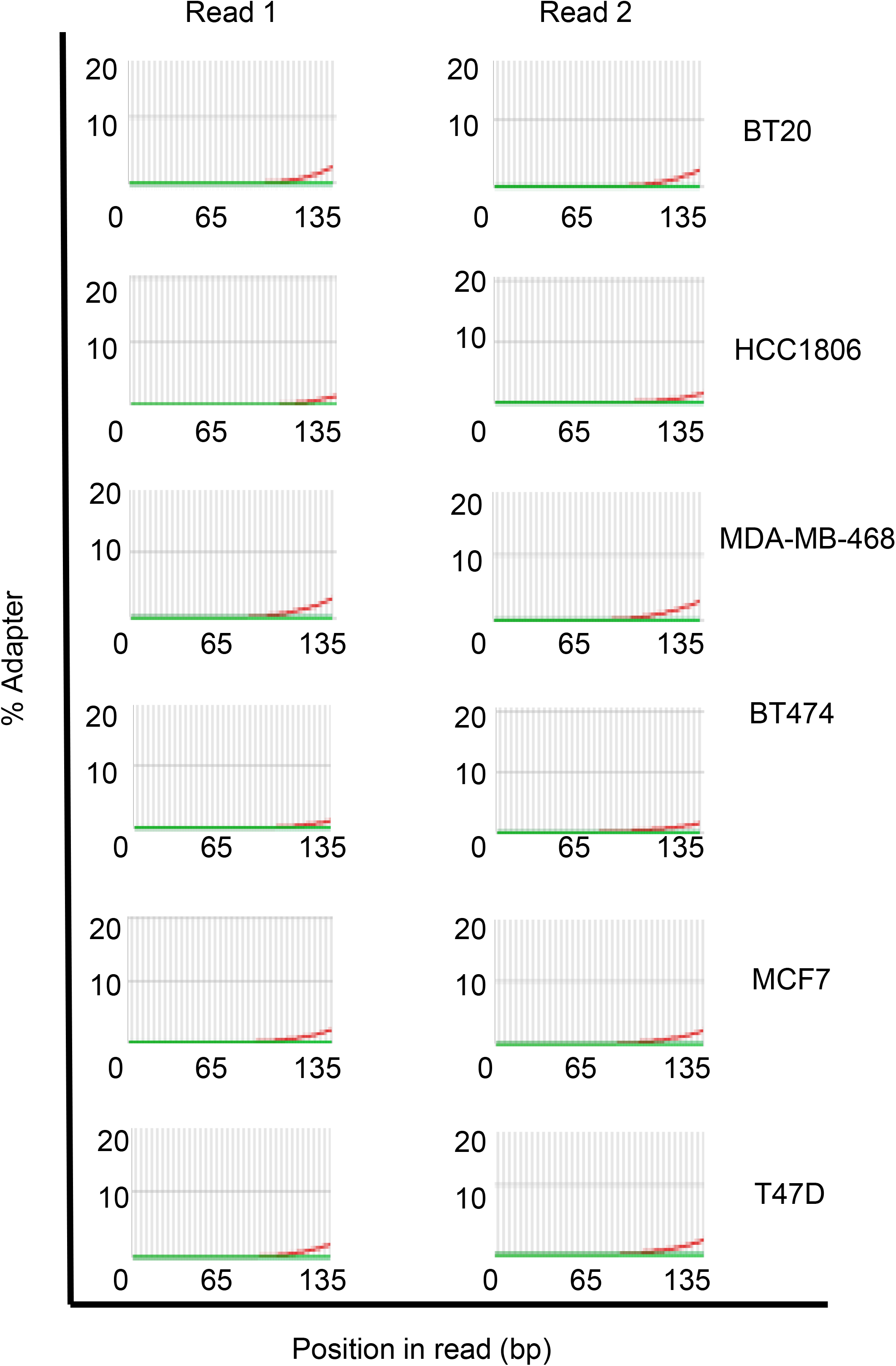
Cumulative presence of adapter sequences. Results show that cumulative presence of adapter sequences is less than 5% in each cell sample, indicating that the data sets could be further analysed without adapter-trimming.

**Fig. S4.**
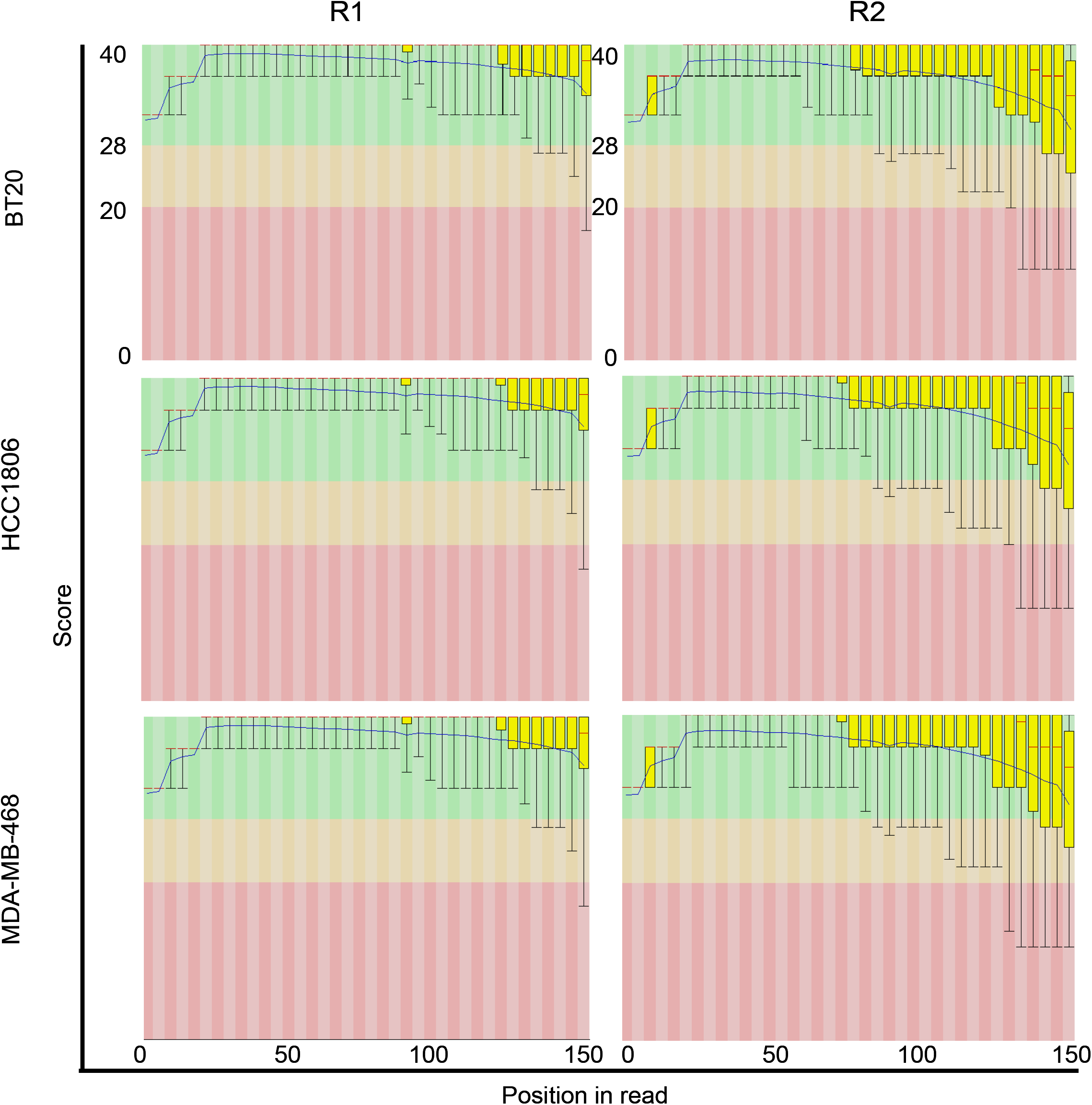
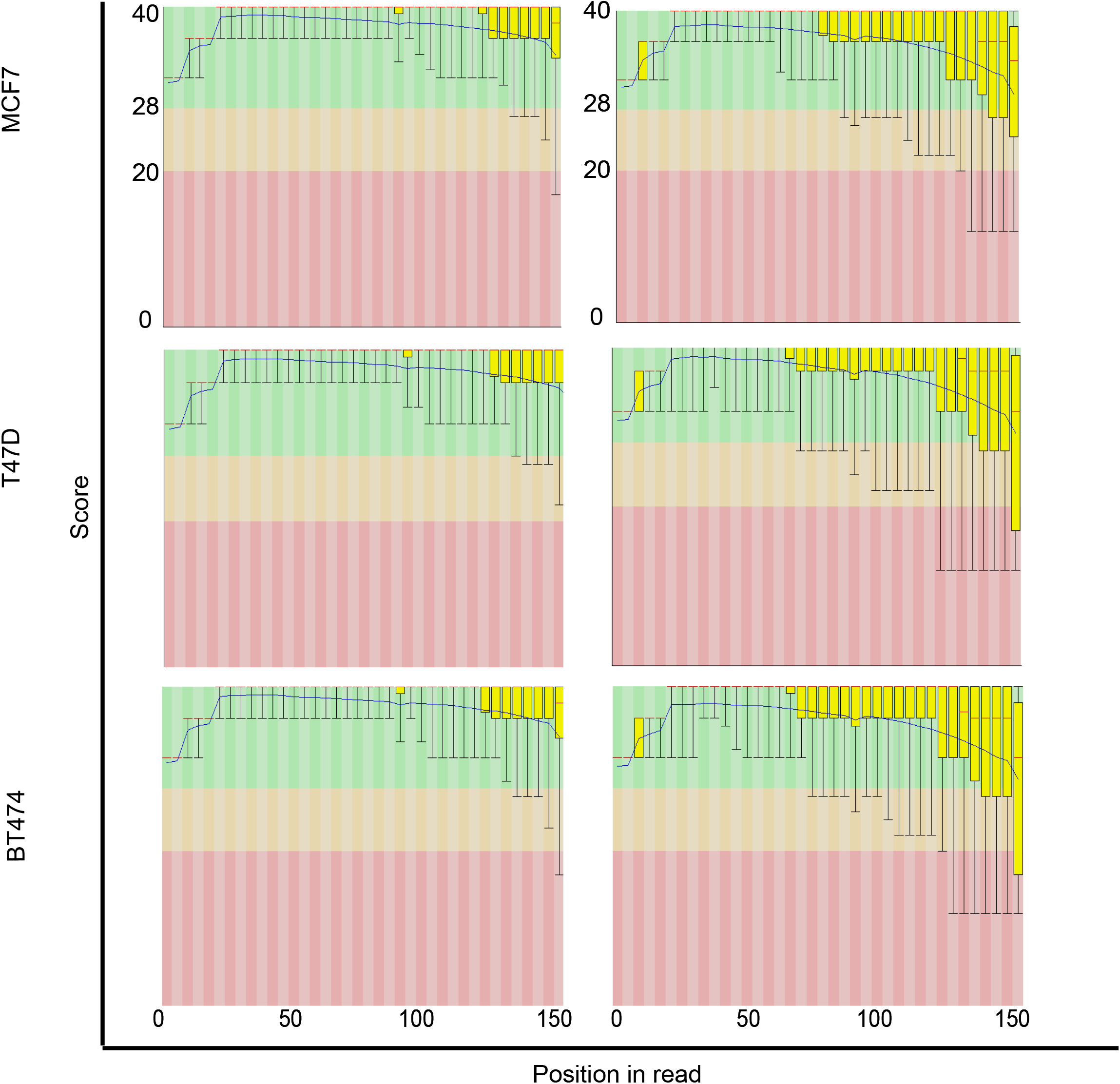
Per base sequence quality for all sequencing data sets. Y axis is divided into good quality calls (coloured green), reasonable quality calls (coloured orange) and poor-quality calls (coloured red). In general, it is normal to observe base calls dropping into the red area towards the end of reads. The blue line representing the mean quality of base calls consistently stayed in the green area, indicating that sequencing data sets were of high quality.

